# Selective disruption of lipid peroxide homeostasis in intratumoral regulatory T cells by targeting FSP1 enhances cancer immunity

**DOI:** 10.1101/2025.07.06.663397

**Authors:** Jesse Garcia Castillo, Stephanie Silveria, Antoine Sauquet, Leo Schirokauer, Joseph Hendricks, Hei Sook Sul, James A. Olzmann, Michel DuPage

## Abstract

A burgeoning approach to treat cancer is the pharmacological induction of ferroptotic cell death of tumor cells. However, the impact of disrupting anti-ferroptotic pathways in the broader tumor microenvironment (TME), such as in immune cells, is still undefined and may complicate treatments. Here, we show that Ferroptosis Suppressor Protein 1 (FSP1*/Aifm2*) is critically required for regulatory T cell (Treg) resistance to ferroptosis and their immunosuppressive function within the TME. Compared to other canonical ferroptosis regulators such as GPX4, GCH1, and NRF2, only FSP1 was induced upon T cell activation. Deletion of *Aifm2* in all T cells, or Tregs specifically, enhanced tumor control by selectively disrupting Treg immunosuppression within tumors without inciting autoimmune pathology in mice. As opposed to deletion of *Gpx4* in all T cells, T cell deletion of *Aifm2* did not impair antigen-specific CD8^+^ T cell responses. These results reveal a unique opportunity for targeting a regulator of ferroptosis that can not only directly target cancer cells, but also simultaneously enhance anti-cancer immune responses without inciting autoimmunity.

## Main Text

Ferroptosis, an iron-dependent form of regulated cell death driven by unchecked lipid peroxide accumulation, has emerged as a promising approach for treating therapy-resistant cancers(*1–3*). Unlike apoptosis, ferroptosis can release danger-associated molecular patterns (DAMPs) that activate immune responses(*4*, *5*). While glutathione peroxidase 4 (GPX4) is the primary defense against ferroptosis, cancer cells often express alternative protective ferroptotic regulators, such as ferroptosis suppressor protein 1 (FSP1) and GTP cyclohydrolase 1 (GCH1), to resist GPX4 inhibition(*6–11*).

Regulatory T cells (Tregs), a suppressive subset of CD4^+^ T cells, pose a major barrier to effective cancer immunotherapy by inhibiting antitumor responses through direct cell contact and the release of immunosuppressive molecules(*12–14*). Tregs are uniquely adapted to the nutrient-deprived and hypoxic tumor microenvironment (TME) by preferentially using oxidative phosphorylation (OXPHOS), importing lipids via CD36, and utilizing lactate and fatty acids as alternative carbon sources for their metabolism(*15–18*). Moreover, Tregs upregulate antioxidant pathways—including glutathione synthesis and thioredoxin—to mitigate oxidative stress and preserve their function in the TME(*19–21*). Although Tregs depend on GPX4 to prevent ferroptosis, inhibiting GPX4 is not a viable strategy for cancer immunotherapy as its inhibition also impairs effector CD8^+^ and CD4^+^ T cell survival(*22*, *23*).

In this study, we establish that FSP1/*Aifm2* is a critical effector protein for Treg resistance to ferroptosis induction *in vitro* and Treg function *in vivo* in tumors. We found that Tregs use FSP1 to prevent lipid peroxide accumulation despite having higher levels of intracellular reactive oxygen species (ROS). *Aifm2*-deficiency was sufficient to increase the sensitivity of Tregs to ferroptosis induction with GPX4 inhibition *in vitro*. *Aifm2*-deficiency in Tregs decreased the expression of proteins essential for Treg effector function, but did not impact effector T cell immune homeostasis or peripheral tolerance in young or aged mice. Total ablation of *Aifm2* in all T cells did not disrupt antitumor T cell responses but did diminish the immunosuppressive function of Tregs. These findings are significant for the translation of pharmacological FSP1 inhibitors for cancer as they show that FSP1 inhibition not only can directly sensitize cancer cells to ferroptosis, but it may also enhance immune responses to cancer by selectively blocking Treg function in the TME without generating autoimmune toxicity.

## Results

### Tregs resist lipid peroxide accumulation and ferroptosis

To understand the regulation of ROS across different T cell populations, we FACS-purified naive Tregs and conventional CD4^+^ T cells (Tconv) for activation *in vitro* and measuring their total ROS and lipid peroxide levels using fluorescent reagents detected by flow cytometry **(Fig. 1A)**. In accordance with numerous studies, we observed that Tregs had increased (∼2-fold) intracellular ROS levels compared to Tconv cells(*18*, *24*) **(Fig. 1B)**. Next, we directly measured lipid peroxidation by measuring the amount of oxidized BODIPY C11 and observed that Tregs had decreased amounts of lipid peroxides compared to Tconv cells **(Fig. 1C)**. While RSL3 increased oxidized C11-BODIPY in both Tregs and Tconv cells, Tconv cells increased CD11-BODIPY^OX^ significantly more than Tregs **(Fig. 1D)**. Importantly, only Ferrostatin-1 (Fer-1), a radical-trapping antioxidant, but not the pan-caspase inhibitor Z-VAD-FMK or RIP1 kinase inhibitor Nectrostatin-1 (Nec-1) treatments reversed CD11-BODIPY^OX^ elevation, as expected **(Fig. 1D)**. Improved control of lipid oxidation ultimately made Tregs more resistant to ferroptotic cell death with the inhibition of GPX4 across a series of concentrations of RSL3 (**Fig. 1E**). Decreased cell viability was confirmed to be due to ferroptosis, as only treatment with Fer-1, but not other programmed cell death inhibitors, was able to rescue cell viability **(Fig. 1F)**. Interestingly, the differences in sensitivity to ferroptosis were not due to the expression of the essential Treg transcription factor Foxp3, as *in vitro* induced Tregs (iTregs), were as sensitive to GPX4 inhibition as Tconv cells (**Fig. 1E**)(*19*). To confirm these findings without activating or culturing T cells long term *in vitro*, we isolated T cells from spleens of mice and treated them with RSL3 alone or RSL3 and Fer-1 for 4 hours *ex vivo*. This revealed that while CD8^+^ and CD4^+^ Tconv cells were highly sensitive to GPX4 inhibition, Tregs remained highly resistant to ferroptosis *ex vivo* **(Fig. 1G)**. Therefore, despite having higher amounts of ROS, Tregs reduce their accumulation of lipid peroxides, which likely increases their resistance to ferroptosis.

**Fig. 1.**
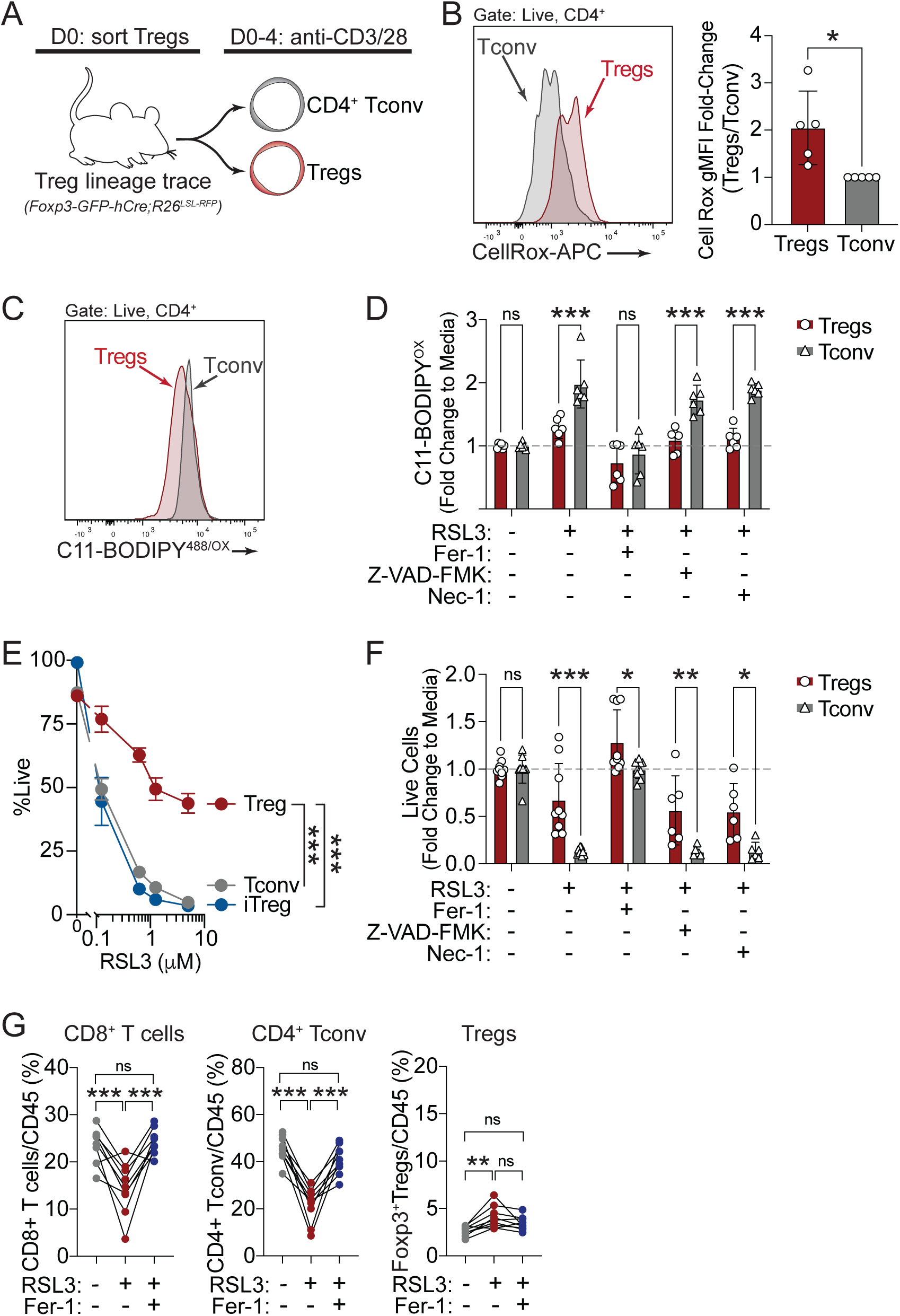
Regulatory T cells (Tregs) naturally resist lipid peroxide accumulation and ferroptosis. (A) Schematic for *in vitro* T cell assays. (B) Representative flow plots (left) and quantification (right) of bulk reactive oxygen species (ROS) by CellROX^TM^ staining in Tregs versus conventional CD4^+^ T cells (Tconv) after activation and culture for 4 days *in vitro*. Data pooled from 5 independent experiments. (C) Representative flow plots for detection of oxidized C11-BODIPY in Tregs versus Tconv that were activated for 4 days *in vitro*. (D) Quantification of oxidized C11-BODIPY in Tregs versus Tconv cells *in vitro* +/- 1μM RSL3 in combination with 2μM Fer-1, 100μM Z-VAD-FMK, or 10μM Nec-1. Data pooled from 3 experiments. (E) Cell viability of Tregs (nTregs), CD4^+^ Tconv, and induced Tregs (Foxp3^+^ iTregs) treated with the indicated concentrations of RSL3 for 4 hours. Representative data from 2 independent experiments. (F) Cell viability of Tregs versus CD4^+^ Tconv treated *in vitro* +/- 1μM RSL3 in combination with 2μM Fer-1, 100μM Z-VAD-FMK, or 10μM Nec-1. Data pooled from 2 experiments. (G) Frequencies of viable CD8^+^ T (left), CD4^+^ Tconv (middle), and Tregs (right) from spleen plotted as the percent of CD45^+^ cells +/- 1μM RSL3 and 2μM Fer-1. Data from n=7 mice. For all plots, **P<0.05, **P<0.01, ***P<0.001 by two-way ANOVA (D-F) or one-way ANOVA (G), or student t-test (B), mean ± s.d (B, D-F) or mean ± s.e.m (G).

### FSP1 expression in Tregs confers resistance to ferroptosis

T cell activation increases levels of ROS, which can be tuned by the expression of antioxidant genes to regulate cytosolic signaling pathways(*22*, *25–29*). Analysis of gene expression by RNA-sequencing of FACS-purified Tregs and Tconv cells that were naive (CD44^lo^CD62L^hi^) versus four days after anti-CD3/anti-CD28 *in vitro* activation, demonstrated that only *Aifm2* increased expression, but not other anti-ferroptotic proteins, e.g. *Gpx4*, *Gch1*, or *Nfe2l2*, upon *in vitro* T cell activation **(Fig. 2A)**(*30*). In addition, a comparison of freshly sorted *in vivo* activated (CD44^hi^CD62L^lo^) Tregs compared to naive Tregs from mice showed a similar induction of *Aifm2* in activated Tregs **(Fig. 2B)**(*30*). Moreover, direct comparison of *in vitro* activated Tregs versus Tconv cells revealed a 2-3 fold greater induction of *Aifm2* and *Gch1* expression, and reduced expression of *Nfe2l2* and *Ascl3*, in Tregs compared to Tconvs **(Fig. 2C)**. These results demonstrate that *Aifm2*/FSP1 is induced during T cell activation, but that Tregs induce higher expression of *Aifm2* and *Gch1* antioxidant genes with activation, which may underlie their enhanced resistance to ferroptosis.

**Fig. 2.**
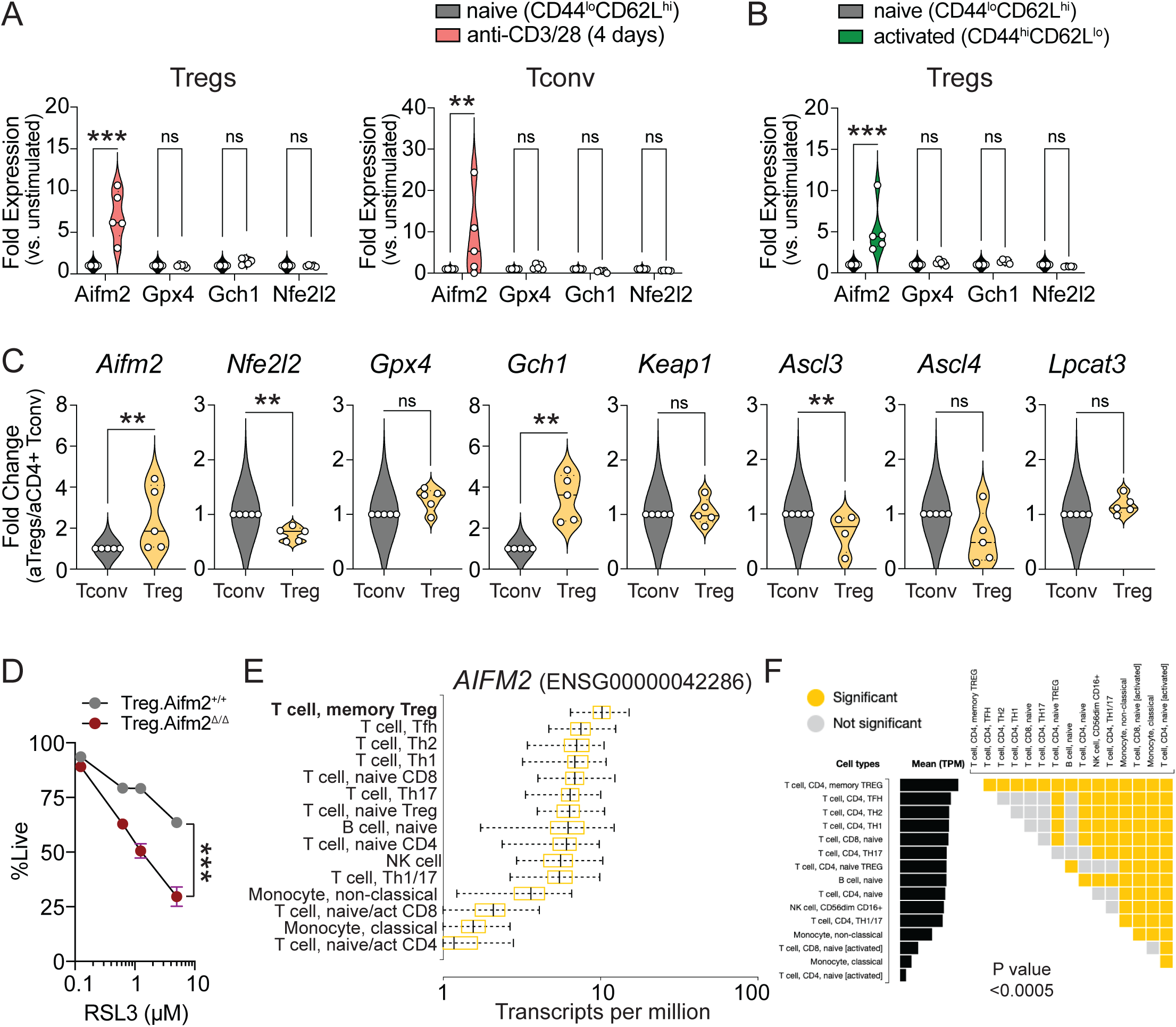
FSP1 drives Treg resistance to ferroptosis. (A) Fold-change expression of indicated anti-ferroptotic genes in naive (CD44^lo^CD62L^hi^) versus in vitro activated (+4 days with anti-CD3/CD28) Tregs. Data from n=5 mice. (B) Fold-change in expression of indicated anti-ferroptotic genes from sorted naive (CD44^lo^CD62L^hi^) versus in vivo activated (CD44^hi^CD62L^lo^) Tregs. (C) Fold change in expression of indicated ferroptosis-related genes in 4 day in vitro activated Treg versus Tconv cells. Data from n=5 mice. (D) Treatment of wildtype versus *Aifm2*-deficient Tregs with different concentrations of RSL3 for 4 hours. (E) *Aifm2* expression from bulk RNA-Seq of human PBMCs populations from Database of Immune Cell eQTLs, expression, epigenomics (DICE). (F) Differential expression analysis of *Aifm2* expression across different PBMC populations. For all plots, **P<0.05, **P<0.01, ***P<0.001 by one-way ANOVA (A-B), student t-test (C), or two-way ANOVA (D), mean ± s.e.m (A-C) or mean ± s.d (D).

While GCH1 was shown to be crucial for T cell proliferative capacity, the role of FSP1 in Treg biology is unexplored(*29*). To define the role of FSP1 in Tregs, we crossed mice carrying LoxP-flanked *Aifm2^fl^*alleles with *Foxp3-GFP-hCre* to specifically delete *Aifm2* in Tregs (called Treg.*Aifm2^Δ/Δ^* mice)(*31*). Treatment of *Aifm2*-deficient Tregs with RSL3 to induce ferroptosis revealed that *Aifm2*-deficient Tregs now exhibited significantly increased sensitivity to RSL3-mediated cell death compared to wildtype Tregs (**Fig. 2D)**. Thus, FSP1 promotes Treg resistance to ferroptosis *in vitro*.

Finally, comparative gene expression analysis from human peripheral blood mononuclear cells (PBMCs) confirmed that human Tregs that are activated (memory Treg) also exhibit the highest expression of *Aifm2* compared both to naive Tregs as well as all other activated T cell populations (**Fig. 2E-F**). These results indicate that FSP1/*Aifm2* expression protects Tregs from ferroptosis and FSP1/*Aifm2* expression is higher in both mouse and human activated Tregs compared to other T cell subsets.

### Treg-specific FSP1 deletion does not impact peripheral tolerance

To directly test whether Treg-specific deletion of *Aifm2* disrupted Treg function *in vivo*, we analyzed lymphoid and nonlymphoid tissues from wildtype versus Treg.*Aifm2^Δ/Δ^* mice at early (16 weeks) and late (28 weeks) stages of life. There were no changes to the frequency of CD4^+^ Tconv, CD8^+^ T, or Tregs across lymphoid organs (lymph node, spleen) at any age (**Fig. 3A and Fig. S1A-B**). However, in the organ tissues examined (colon, skin, lung), while CD4^+^ Tconv and CD8^+^ T cell frequencies did not change in Treg.*Aifm2^Δ/Δ^* mice, we did note a modest reduction in Treg frequency in the lung and colon of Treg.*Aifm2^Δ/Δ^*mice **(Fig. 3A and Fig. S1C)**. Additionally, there were no changes to the activation state of CD4^+^ Tconv, CD8^+^ T, or Tregs in all tissues examined (**Fig. 3B and Fig. S1D**). These data suggest that Treg-specific *Aifm2*-deficiency does not increase the prevalence of autoimmunity in mice.

**Fig. 3.**
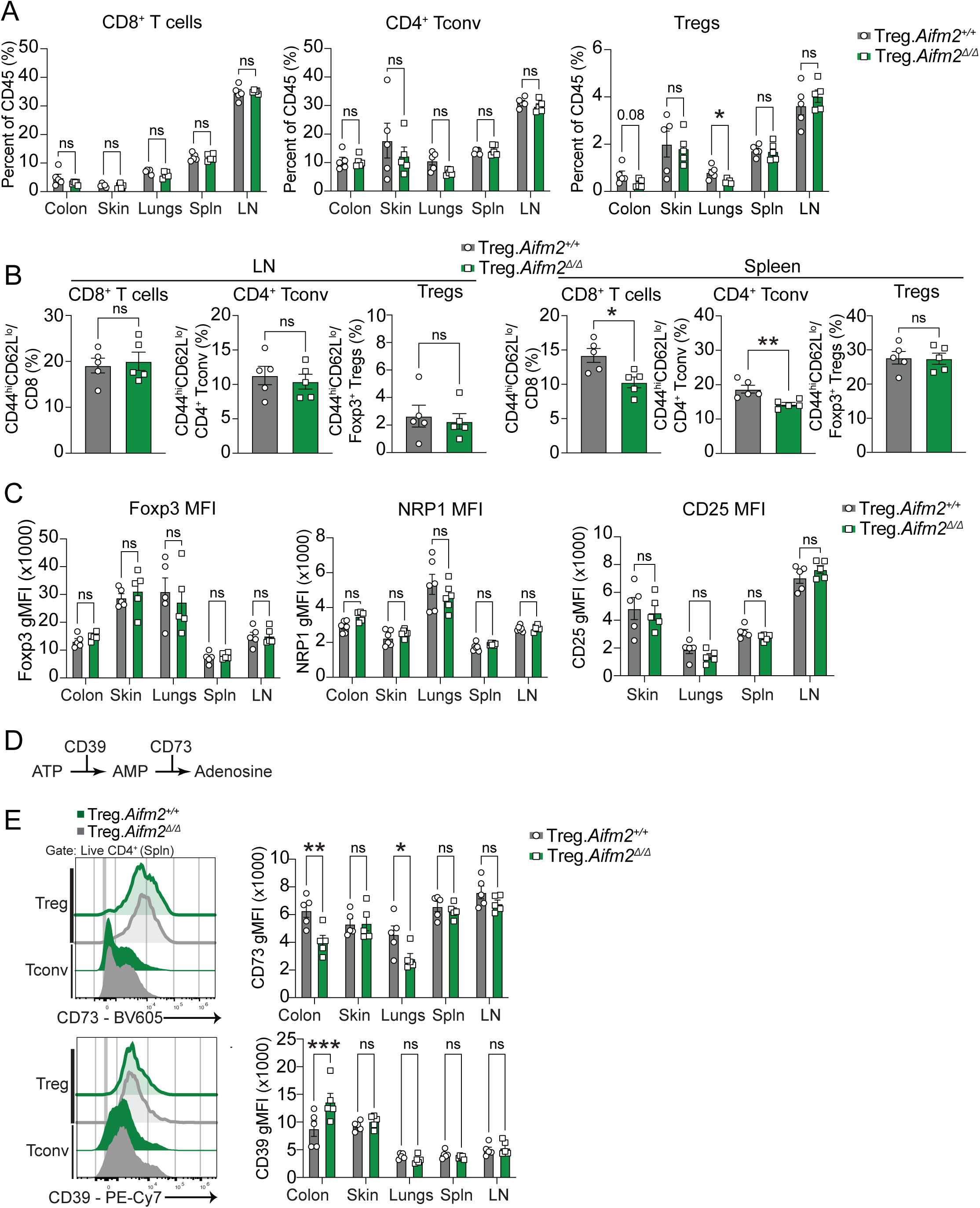
Deletion of *Aifm2* in Tregs does not disrupt peripheral tolerance in mice. (A) Frequency of total CD8^+^ T cells (left), CD4^+^ Tconv cells (middle), and Tregs(right) as a percentage of total CD45^+^ immune cells in lymphoid tissues and organ tissues in 16 week mice. (B) Frequencies of activated (CD44^hi^CD62^lo^) CD8^+^ T cells (left), CD4^+^ Tconv cells (middle), and Tregs (right) in secondary lymphoid organs (SLOs) in 16 week mice. (C) gMFI of Foxp3, CD25, and NRP1 in Foxp3 Tregs across lymphoid tissues and non-lymphoid tissues in 16 week mice. (D) Schematic of pathway for conversion of ATP to Adenosine mediated by CD39 and CD73. (E) Representative histograms and gMFI of CD73 and CD39 in wildtype versus *Aifm2*-deficient Tregs across lymphoid tissues and nonlymphoid tissues in 16 week mice. Data from n=5 for Treg.*Aifm2*^+/+^ and n=6 for Treg.*Aifm2*^Δ/Δ^ mice for 16 week mice. For all plots, **P<0.05, **P<0.01, ***P<0.001 by two-way ANOVA (A, C, E) or student’s t-test (B), mean ± s.e.m.

Phenotyping of the Tregs from Treg.*Aifm2^Δ/Δ^* mice revealed no significant changes in the expression of canonical lineage defining Treg markers (i.e. CD25, Foxp3, and NRP1) in either lymphoid or organ tissues **(Fig. 3C and Fig. S1E)**. However, there was a significant decrease in CD73 expression on colonic and lung Tregs **(Fig. 3D-E).** CD73, in conjunction with CD39, acts as an ectonucleoside that breaks down ATP to generate adenosine, a critical immunosuppressive molecule generated by Tregs. Interestingly, both CD73 and CD39 expression were significantly decreased in the lymphoid tissues of aged mice **(Fig. S1E)**. Altogether, these results show that *Aifm2*-deficiency in Tregs does not dramatically impact the frequencies or phenotype of Tregs in lymphoid or non-lymphoid tissues. Thus, *Aifm2*-deficiency in Tregs does not lead to a systemic breakdown in immunotolerance or organ-specific autoimmunity.

### FSP1-deficiency specifically in Tregs impairs immunosuppression in response to cancer

Since the pharmacological inhibition of FSP1 in cancer cells is moving to clinical trials, we next wanted to assess tumor Treg responses to the inhibition of FSP1. First, we tested whether different subsets of T cells from tumors exhibited different levels of resistance to ferroptosis induction, similar to experiments in Figure 1G. Here, we grew MC38 colon carcinoma tumors subcutaneously for 14 days in wildtype mice, then harvested tumors, made single cell suspensions, and treated all cells for four hours with or without 1μM RSL3 to induce ferroptosis. Strikingly, Tregs again exhibited strong resistance to the induction of ferroptosis with GPX4 inhibition as compared to Tconv and CD8^+^ T cells **(Fig. 4A)**. To test whether Treg-specific FSP1 activity promoted tumor Treg resistance to ferroptosis and supported tumor growth, we compared MC38 tumor growth in wildtype versus Treg.*Aifm2^Δ/Δ^* mice **(Fig. 4B)**(*32*). Tumor growth was significantly reduced in Treg.*Aifm2^Δ/Δ^*mice **(Fig. 4C)**. While Treg.*Aifm2^Δ/Δ^* mice did not exhibit changes to Treg frequencies or tumor-specific CD8^+^ T cells in tumors (**Fig. 4D and Fig. S2A),** Treg.*Aifm2^Δ/Δ^* mice did exhibit increased CD8 to Treg ratios in tumors, indicative of a selective impact on Tregs compared to effector T cells in tumors with FSP1-deficiency in Tregs (**Fig. 4D)**. In addition, we did not observe altered expression of the exhaustion markers PD-1 or LAG-3 in CD8^+^ T cells in Treg.*Aifm2^Δ/Δ^* mice (**Fig. S2B**). Importantly, to demonstrate that the reduction in tumor growth was due to the accumulation of lipid peroxides in FSP1-deficient Tregs, we treated wildtype and Treg.*Aifm2^Δ/Δ^* tumor-bearing mice with the radical trapping antioxidant liproxstatin-1 (Lip-1), a ferroptosis inhibitor that is functional *in vivo in mice*(*33*) (**Fig. 4E**). Lip-1 treatment restored tumor growth in Treg.*Aifm2^Δ/Δ^* mice, while Lip-1 treatment had no effect on tumor growth in wildtype mice (**Fig. 4F**). These results indicate that FSP1 activity in Tregs is required to control their accumulation of lipid peroxides within the TME, and disruption of FSP1 activity can release immunosuppression by Tregs and promote better tumor control.

**Fig. 4.**
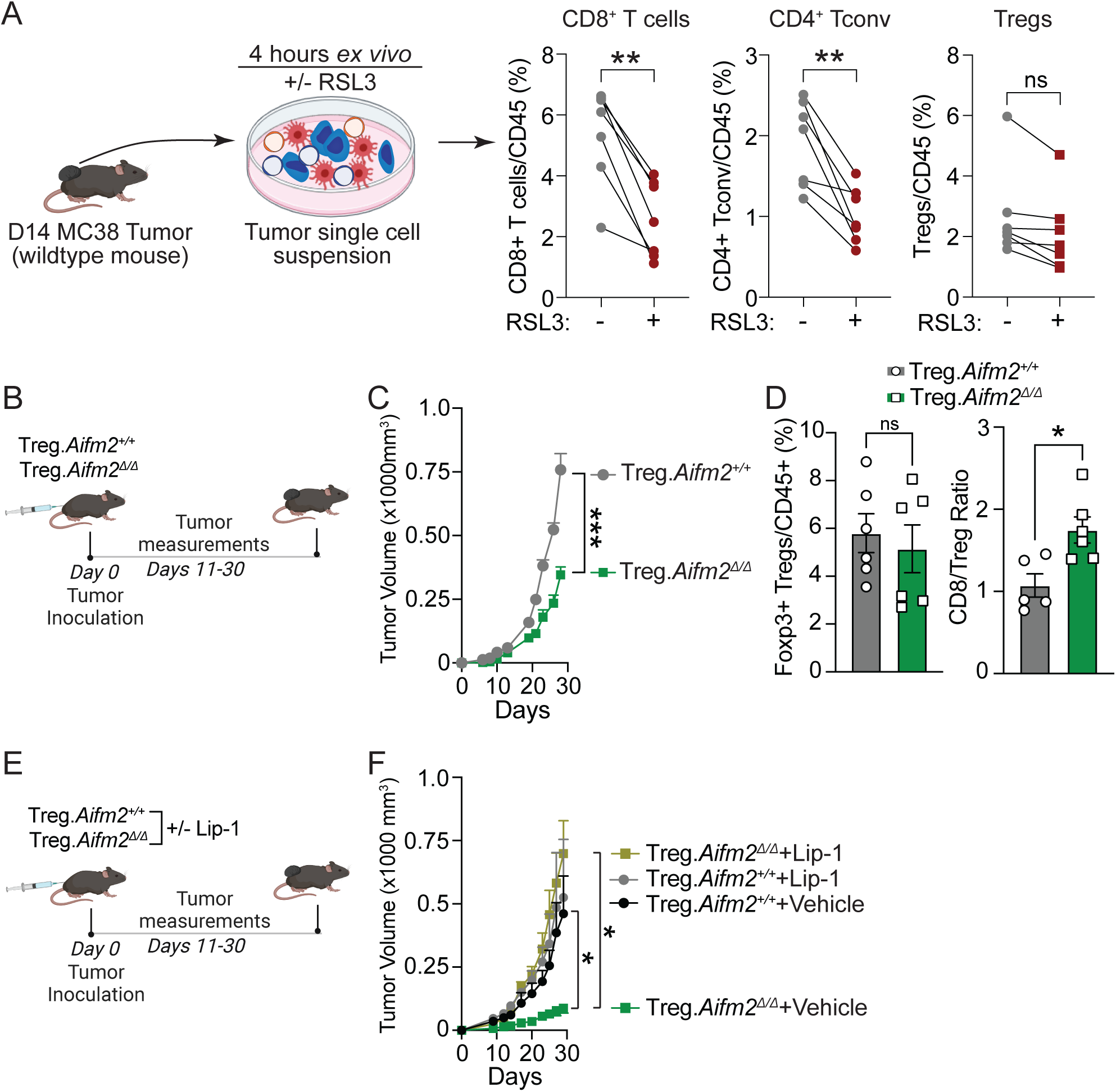
Deletion of *Aifm2* in Tregs improves cancer immunity. (A) Schematic for treating single cell suspensions from MC38 tumors +/- 1μM RSL3 (left). Frequencies of viable CD8^+^ (left), CD4^+^ Tconv (middle), and Tregs (right) from tumor single cell suspensions as a percentage of CD45^+^ cells +/- 1μM RSL3 for 4 hours. Data from n=6 mice. (B) Experimental schematic for measuring tumor growth in wildtype versus mice with *Aifm2*-deficient Tregs. (C) Growth of MC38 tumors in wildtype versus mice with *Aifm2*-deficient Tregs. Data from n=5 for Treg.Aifm2^+/+^ and n=6 from *Treg.Aifm2^Δ/Δ^*. (D) Frequency of Foxp3^+^ Tregs as a percentage of total CD45^+^ immune cells (left) and CD8:Treg ratio (right) in MC38 tumors at D29. (E) Experimental schematic for measuring tumor growth in wildtype versus mice with *Aifm2*-deficient Tregs with or without 10mg/kg liproxstatin injected i.p daily. Data from n=3 for Treg.*Aifm2^+/+^* + Vehicle, n=4 for Treg.*Aifm2^+/+^* + Lip-1, n=5 for Treg.*Aifm2^Δ/Δ^*+ Vehicle, and n=5 for Treg.*Aifm2^Δ/Δ^* + Lip-1. (F) Growth of MC38 tumors in wildtype versus mice with *Aifm2*-deficient Tregs +/- Lip-1. For all plots, **P<0.05, **P<0.01, ***P<0.001 by student t-test (A, D) or two-way ANOVA (C, F), mean ± s.e.m.

### FSP1-deficiency in all T cells enhances T cell responses to infection and cancer

T cell deficiency for *Gpx4*- or *Gch-1* was recently shown to negatively impact T cell responses in several disease settings(*23*, *29*, *34*). To test if *Aifm2*-deficiency also impacts all T cell responses, which would occur with pharmacological inhibition for cancer treatment, we generated *Aifm2*-deficient T cell mice (*CD4-Cre;Aifm2^fl/fl^*, called *T.Aifm2^Δ/Δ^*) and infected these mice with *Listeria monocytogenes*, an intracellular pathogen known for stimulating a robust CD8^+^ T cell response (**Fig. S3A**)(*35*, *36*). Importantly, *T.Aifm2^Δ/Δ^*mice had no deficiency in bacterial control, as there was no difference in bacterial CFUs recovered from spleens or livers of wildtype versus *T.Aifm2^Δ/Δ^*six days post infection (**Fig. S3B**). Upon *Listeria* infection, we observed a decrease in the frequency of Tregs as a percentage of CD4^+^ T conv cells, as well as of total CD45^+^ immune cells, in *T.Aifm2^Δ/Δ^* (**Fig. S3C**). The frequency of total CD8^+^ T cells, *Listeria*-specific CD8^+^ T cells, or CD4^+^ Tconv cell populations, however, did not change (**Fig. S3D-F**).

Next, to test the impact of deleting *Aifm2* in all T cells, as would happen with a pharmacological FSP1 inhibitor, in the setting of cancer, we implanted MC38 tumors into wildtype versus *T.Aifm2^Δ/Δ^* mice and measured tumor growth (**Fig. 5A**). Tumor growth was significantly decreased in mice harboring *Aifm2*-deficient T cells (**Fig. 5B**). While there was no change in the frequency of Tregs in MC38 tumors, there was an increase in the CD8:Treg ratios in tumors from *T.Aifm2^Δ/Δ^* compared to wildtype mice, as observed with Treg-specific *Aifm2*-deficiency (**Fig. 5C-D**)(*37*, *38*). Assessment of CD8^+^ T cell exhaustion in tumors by staining for both PD-1 and Lag-3 proteins showed no changes in total CD8^+^ T cells **(Fig. 5E)**. However, tumor-specific CD8^+^ T cells, while unchanged in total frequencies, exhibited decreased frequencies of PD-1^+^Lag3^+^ MuLV/H2K^b+^ CD8^+^ T cells (**Fig. 5F**)(*39*). These results starkly contrast with the consequences of T cell deficiencies in other antioxidant genes, i.e. *Gpx4* or *Gch1*, where deficient T cells completely lost their capacities to control infections or cancer(*23*, *29*). Instead, deficiency for FSP1/*Aifm2* in all T cells completely preserves CD8^+^ and CD4^+^ Tconv cell function while potently disrupting the function of Tregs.

**Fig. 5.**
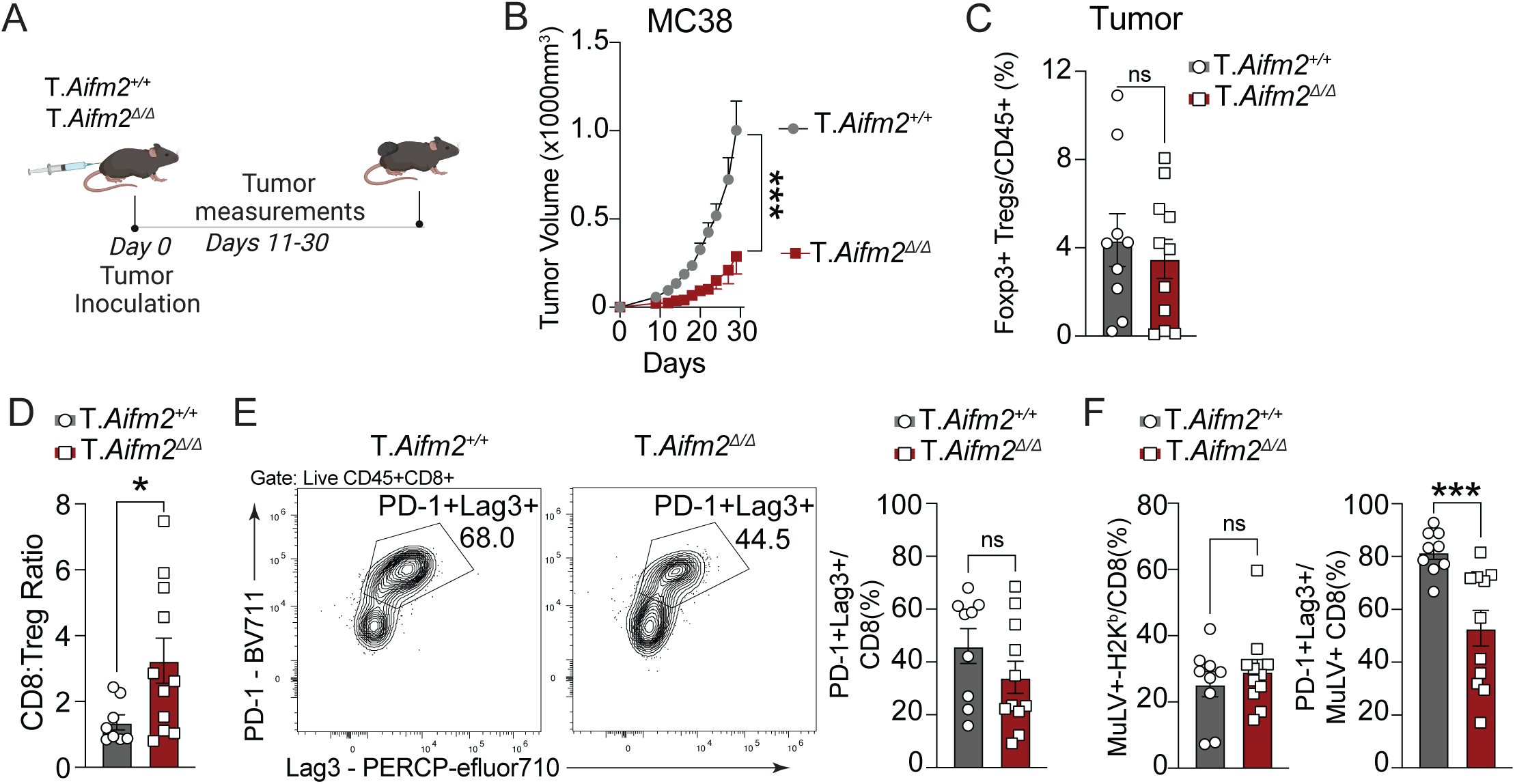
Total T cell deficiency for *Aifm2* preserves CD8^+^ T cell function and improves tumor immunity. (A) Experimental schematic for measuring tumor growth in wildtype versus mice with *Aifm2*-deficient T cells. (B) Growth of MC38 tumors in wildtype versus mice with *Aifm2*-deficient T cells. Data shown from n=5-6 mice/group from one of two experiments. (C) Frequency of Foxp3^+^ Tregs as a percentage total CD45^+^ immune cells in MC38 tumors at D29 from (B). (D) CD8:Treg ratio in MC38 tumors at D29 from (B). (E) Representative flow plot for PD-1 and LAG3 staining in total CD8^+^ T cells (left) and quantification of the frequency of PD-1^+^Lag3^+^ CD8^+^ T cells in MC38 tumors (right). (F) Frequency of tumor-specific (MuLV/H2K^b+^) as a percentage of CD8^+^ T cells (left) and frequency of PD-1^+^Lag3^+^ of MuLV/H2K^b+^ CD8^+^ T cells in MC38 tumors (right). For all plots, **P<0.05, **P<0.01, ***P<0.001 by two-way ANOVA (B) or student t-test (C, D, F), mean ± s.e.m.

## Discussion

In this study, we demonstrate that FSP1 (*Aifm2*) plays a critical role, and unique role amongst T cells, in sustaining Treg function within tumors. Unlike other ferroptosis resistance factors such as GPX4, which are essential for proper T cell function during infections or cancer, FSP1 appears to be selectively required for Tregs without compromising effector T cell responses. We show that FSP1 expression is induced during T cell activation, distinguishing it from other ferroptosis regulators like GPX4, GCH1, or NRF2. Our data indicate that FSP1 supports Treg survival and suppressive function by limiting lipid peroxide accumulation despite high levels of ROS generation in Tregs. This suggests that increased FSP1 expression in Tregs may have evolved to prevent sublethal oxidative damage that would otherwise kill or impair Tregs.

These findings reveal a novel paradigm in which targeting FSP1 may specifically modulate Treg-mediated immunosuppression in cancer without affecting effector T cells. In contrast to GPX4-targeted approaches, which can dampen overall T cell responses and carry a narrow therapeutic window due to GPX4’s essential role in multiple tissues, inhibition of FSP1 appears to selectively compromise Treg function(*23*, *34*, *40*, *41*). In addition, Tregs are sensitive to specific antioxidant pathway activity as the genetic deletion of KEAP1 leads to loss of Treg lineage identity that leads to spontaneous autoimmunity due to hyperactivation of NRF2 and uncontrolled induction of antioxidant pathways(*42*). Our tumor models showed that genetic ablation of FSP1 in T cells resulted in better tumor control, potentially due to a reduction in Treg suppressive capacity and a concomitant decrease in canonical exhaustion markers (e.g., PD-1, Lag3) on tumor-specific CD8^+^ T cells(*43*).

The implications of these findings are significant. By specifically targeting FSP1, it may be possible to alleviate the immunosuppressive effects of Tregs within the TME, thereby enhancing the efficacy of cancer immunotherapies. Moreover, FSP1 inhibition exhibits great potential to synergize with immune checkpoint blockade (e.g., anti-PD-1 therapies) and cellular immunotherapies such as CAR T cells by preventing T cell exhaustion and preserving cytotoxic function. PD-1 blockade has also been shown to induce IFN-γ production by CD8^+^ T cells, which induced ferroptosis in cancer cells directly by impacting the lipid composition through Ascl4 expression(*44*). Thus, this dual approach, targeting tumor cells directly through ferroptosis induction while simultaneously modulating Treg function, could offer a much broader therapeutic strategy for treating cancer.

Our study also underscores the importance of metabolic regulation in Treg biology. Tregs exhibit a unique metabolic profile characterized by a reliance on oxidative phosphorylation and fatty acid utilization, which supports their survival in the nutrient-deprived, hypoxic TME(*15–17*). Despite high ROS levels due to these forms of metabolism, Tregs maintain low lipid peroxide accumulation, in part by maintaining higher expression of FSP1. Disrupting this balance through FSP1 inhibition leads to functional defects in Tregs without appearing to cause their complete elimination in vivo, which is preferable for preserving immune homeostasis.

In conclusion, our results position FSP1 as a Treg-specific ferroptosis regulator and a promising therapeutic target for enhancing cancer immunity. By selectively modulating FSP1 activity, it may be possible to reduce Treg-mediated suppression in tumors, preserve effector T cell responses, and ultimately improve the outcomes of cancer immunotherapy. Future studies should focus on translating these findings into pharmacological interventions, as well as exploring the potential of FSP1-targeting strategies in combination with existing immunotherapies to achieve synergistic effects.

## Materials and Methods

### Mice

CD4-Cre mice were obtained from Jackson laboratories (JAX:022071) and bred in house. Foxp3-GFP-hCre mice were obtained from Jackson laboratories (JAX:023161) and bred in house. *Aifm2^fl^* mice were generated as described(*31*). For tumor studies, syngeneic C57BL/6J mice were inoculated with 5.0×10^5^ MC38 in PBS subcutaneously. Tumor measurements were performed blindly across the entire experiment by a single operator measuring three dimensions of the tumor with calipers three times per week. For in vivo liproxstatin treatments (Selleck Chemicals), mice were injected intraperitoneally with 10mg/kg of liproxstatin-1 dissolved in DMSO and diluted in PEG300, Tween-80, and dH2O. All the experiments were conducted according to the Institutional Animal Care and Use Committee guidelines of the University of California, Berkeley.

### Cell lines

MC38 cell lines were kindly provided by Dr. Jeff Bluestone’s lab(*45*). All cell lines were maintained in DMEM (GIBCO) supplemented with 10% FBS, sodium pyruvate (GIBCO), 10mM HEPES (GIBCO), and penicillin-streptomycin (GIBCO). Tumor cells were grown at 37℃ with 5% CO_2_.

### Listeria monocytogenes strains

All strains of L. monocytogenes were derived from the wild-type 10403S strain. All strains were cultured in filter-sterilized nutrient-rich Brain Heart Infusion (BHI) media (BD Biosciences) containing 200 μg/mL streptomycin (Sigma-Aldrich).

### Intravenous and Intratumoral Listeria infection

Overnight cultures were grown in BHI + 200 μg/mL streptomycin at 30℃. The following day, bacteria were grown to logarithmic phase by diluting the overnight culture in fresh BHI + 200 μg/mL streptomycin and culturing at 37℃ shaking. Log-phase bacteria were washed and frozen in 9% glycerol/PBS. For intravenous infections, frozen stocks were diluted in PBS to infect via the tail vein with 5×10^3^ CFU log-phase bacteria. The mice were euthanized 6 days after intravenous injections and organs were collected for CFU or flow analysis.

### Primary Treg and T cell culture

Spleens and lymph nodes were collected from Foxp3-GFP-hCre;R26^LSL-RFP^; *Aifm2^+/+^*, Foxp3-GFP-hCre;R26^LSL-RFP^; *Aifm2^Δ/Δ^*, CD4-Cre;*Aifm2^+/+^* or CD4-Cre;*Aifm2^Δ/Δ^*. Single cell suspensions were generated and enriched for CD4+ T cells by negative selection using EasySep magnetic bead kit (STEMCELL Technologies) and stained with anti-mouse CD4 (RM4-5, BioLegend), anti-CD62L (MEL1-14, Biolegend), anti-CD8 (53-6.7, Biolegend), anti-CD25 (PC61, BioLegend), anti-CD357 (GITR) (DTA-1, Biolegend). Naïve Treg cells (CD4^+^Foxp3GFP^+^RFP^+^CD62L^+^ or CD4^+^CD25^+^GITR^+^CD62L^+^) and naive CD4^+^ T cells (CD4^+^CD62L^+^Foxp3GFP^-^RFP^-^ or CD4^+^CD62L^+^GITR^-^CD25^-^) were sorted using an Aria Fusion sorter (BD Biosciences) with a 70μm nozzle. Cells were activated with anti-CD3 and anti-CD28 coated beads (Dynabeads Mouse T-Activator CD3/CD28, Invitrogen) at a ratio of 1:1 for Tconv or 1:3 ratio for Tregs (cell:bead). Cells were kept at a concentration of 10^6^ cells/ml in DMEM medium supplemented with 10% FBS, non-essential amino acids, sodium pyruvate, L-glutamine, HEPES, and β-ME, and cells cultured more than 24 hr were supplemented with 200–2,000 IU/ml recombinant human IL-2 (NIH).

### Tissue Collection and preparation for Flow cytometry

Flow cytometry was performed on an BD LSR Fortessa X20 (BD Biosciences), CyTEK Aurora (CyTEK Biosciences) or LSRFortessa (BD Biosciences) and datasets were analyzed using FlowJo software (Tree Star). Single cell suspensions were prepared in ice-cold FACS buffer (PBS with 2mM EDTA and 1% BS) and subjected to red blood cell lysis using ACK buffer (150mM NH_4_Cl, 10mM KHCO_3_, 0.1mM Na2EDTA, pH7.3). Dead cells were stained with Live/Dead Fixable Blue or Aqua Dead Cell Stain kit (Molecular Probes) in PBS at 4℃. Cell surface antigens were stained at 4℃ using a mixture of fluorophore-conjugated antibodies. Surface marker stains for murine samples were carried out with anti-mouse CD3 (17A2, BioLegend), anti-mouse CD4 (RM4-5, BioLegend), anti-mouse CD8a (53-6.7, BioLegend), anti-mouse, CD44 (IM7, BioLeged), anti-mouse CD45 (30-F11, BioLegend), anti-H-2K^b^ MuLV p15E Tetramer-KSPWFTTL (MBL), anti-H-2K^b^ SIINFEKL Tetramer (NIH Tetramer Core) in PBS, 0.5% BSA. Cells were fixed using the eBioscience Foxp3/Transcription Factor staining buffer set (eBioscience), prior to intracellular staining. Intracellular staining was performed using anti-mouse Foxp3 (FJK-16S, eBioscience) at 4℃, according to manufacturer’s instructions. Cells were resuspended in PBS and filtered through a 70-μm nylon mesh before data acquisition. Datasets were analyzed using FlowJo software (Tree Star).

### Statistical Methods

p values were obtained from unpaired two-tailed Student’s t tests for all statistical comparisons between two groups, and data were displayed as mean 土 SEMs or mean土 SD. For multiple comparisons, one-way ANOVA or two-way ANOVA was used. For tumor growth curves, two-way ANOVA was used with Sidak’s multiple comparisons test performed at each time point or by multiple regression analysis p values are denoted in figures by *p < 0.05, **p < 0.01, ***p < 0.001, and ****p < 0.0001.

## Acknowledgments

We thank Hector Nolla, Alma Valleros, Kartoosh Heydari and Anita Wong Lin of the UC Berkeley Cancer Research Laboratory Flow Cytometry Facility. This research was supported by 1DP2CA247830-01 from the National Institutes of Health (NCI), R01GM112948 from the National Institutes of Health, and C23CR5612 from the UC CRCC Cancer Research Coordinating Committee. M.D. is a Pew-Stewart Scholar and a St. Baldrick’s Scholar with generous support from Hope with Hazel. J.G.C is a HHMI Gilliam Fellow. J.A.O is a Chan-Zuckerberg investigator.

## Author Contributions

Conceptualization and Methodology, M.D., J.A.O., J.G.C.; Investigation, J.G.C., S.S., A.S., L.S., J.H., J.A.O., and M.D.; Writing – Original Draft, J.G.C., M.D., J.A.O.; Writing – Review & Editing, J.G.C., M.D., J.A.O., S.S.; Supervision, M.D., J.A.O., J.G.C.; Funding Acquisition, M.D., J.A.O.

## Resource Availability

### Lead Contact

Further information and requests for resources and reagents should be directed to and will be fulfilled by the lead contact, Michel DuPage.

### Materials availability

*Listeria* monocytogenes strains used in the study will be made available from the lead contact upon request.

## Supplementary Materials

Materials and Methods

Figures S1 to S3

