## Supplemental Figures S1-3 for "Selective disruption of lipid peroxide homeostasis in intratumoral regulatory T cells by targeting FSP1 enhances cancer immunity"

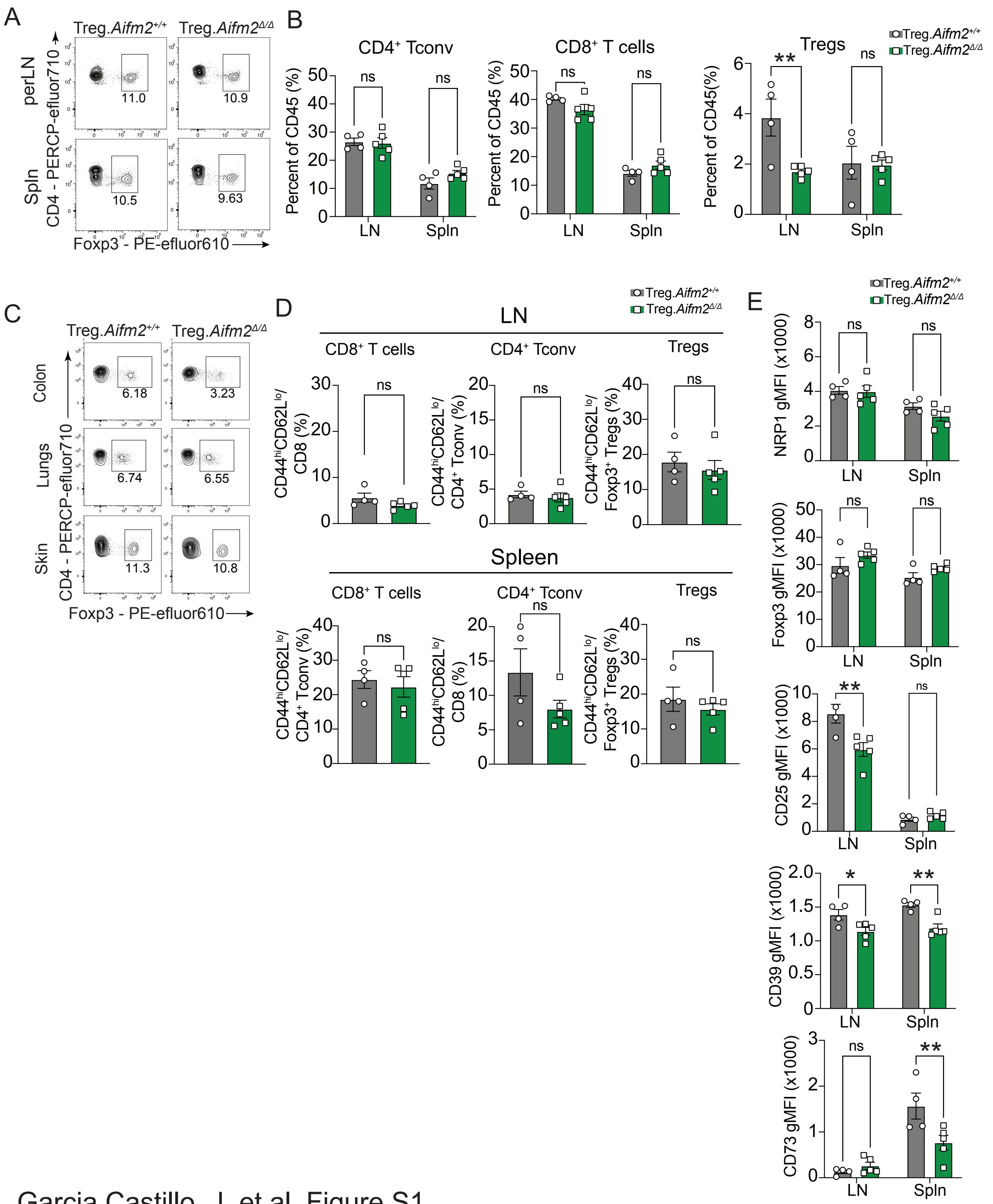


Garcia Castillo, J. et al. Figure S1.

**Fig. S1 | Deletion of *Aifm2* differentially does not impact the frequency of T cell populations across different tissues**. (A) Representative flow plots for wildtype versus *Aifm2*-deficient Tregs from peripheral lymph nodes (LN) and spleen. (B) Frequency of bulk $CD8^+$ T cells (left), $CD4^+$ Tconv cells (middle), and Tregs (right) as a percentage of total $CD45^+$ immune cells in colon, lungs, and skin across lymphoid tissues and nonlymphoid tissues in 28 week mice. (C) Representative flow plots for wildtype versus *Aifm2*-deficient Tregs from peripheral colon, skin, and lungs. (D) Frequency of $Foxp3^+$ Tregs as a percentage of total $CD45^+$ immune cells in lymph nodes and spleens in 28 week mice. (E) gMFI of NRP1, Foxp3, CD25, CD39, and CD73 in wildtype versus *Aifm2*-deficient Tregs across lymphoid tissues of 28 week mice. Data from n=4 for Treg.$Aifm2^{+/+}$ and n=5 for Treg.$Aifm2^{\Delta/\Delta}$ mice for 28 week mice. For all plots, **P<0.05, **P<0.01, ***P<0.001 by two-way ANOVA (B, D, E), mean ± s.e.m.

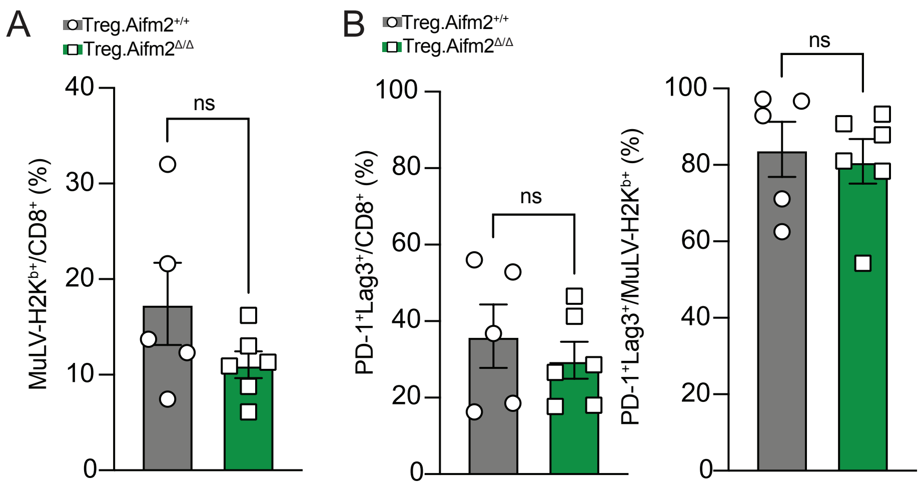


Garcia Castillo, J. et al. Figure S2.

**Fig. S2 | No changes in tumor-specific CD8+ T cell frequency and phenotype in tumors from mice with Treg-specific *Aifm2* deficiency**. (A) Frequency of tumor-specific (MuLV/H2Kb$^{+}$) as a percentage of CD8$^{+}$ T cells in MC38 tumors from Treg.*Aifm2*$^{+/+}$ or Treg.*Aifm2*$^{\Delta/\Delta}$ mice. (B) Frequency of PD-1$^{+}$Lag3$^{+}$ of bulk CD8$^{+}$ T cells (right) or MuLV/H2Kb$^{+}$ CD8$^{+}$ T cells (left) in MC38 tumors from Treg.*Aifm2*$^{+/+}$ or Treg.*Aifm2*$^{\Delta/\Delta}$ mice. Data from n=5 for Treg.*Aifm2*$^{+/+}$ and n=6 for Treg.*Aifm2*$^{\Delta/\Delta}$ mice. For all plots, **P<0.05, **P<0.01, ***P<0.001 by student t-test, mean ± s.e.m.

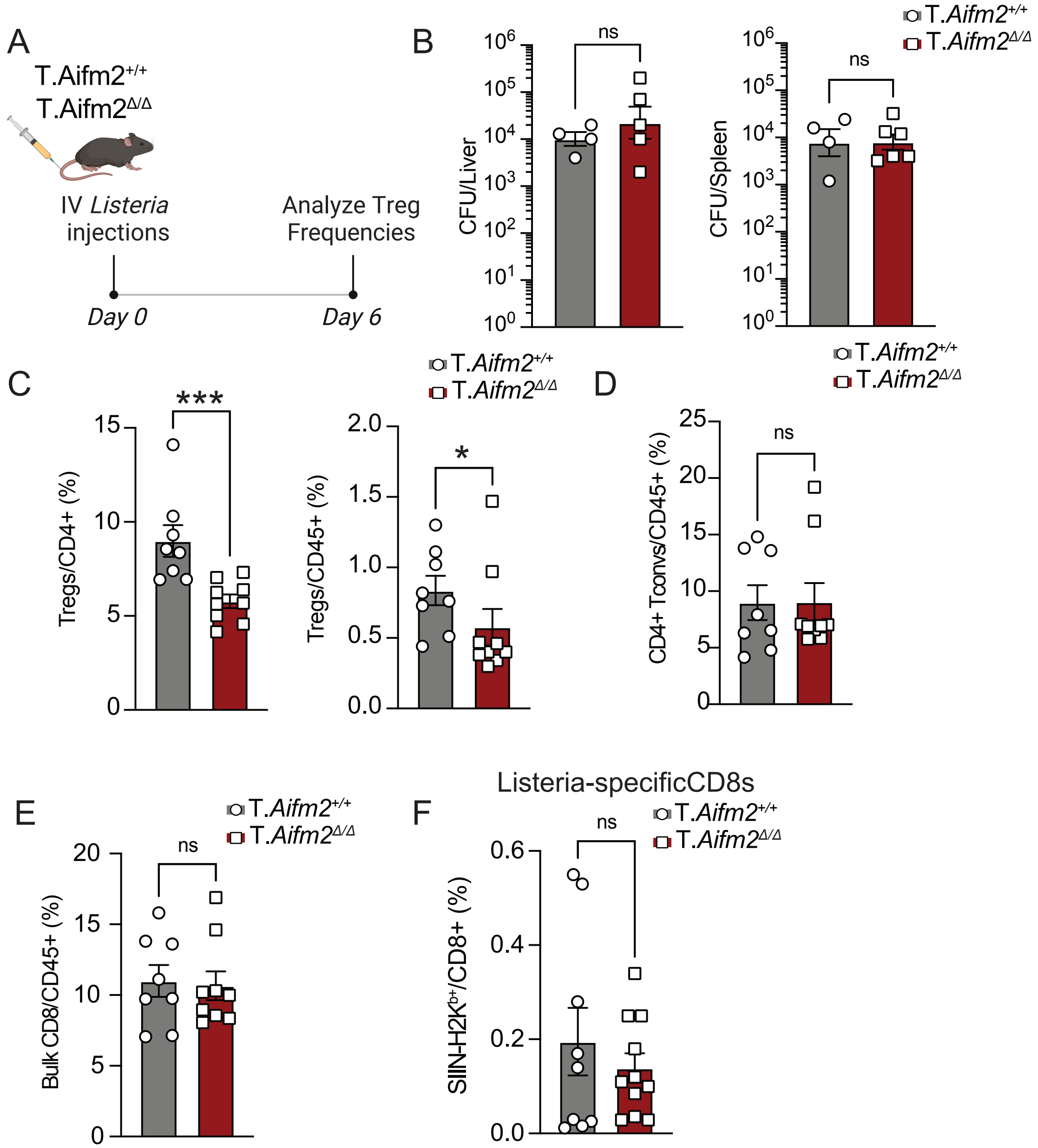


Garcia Castillo, J. Figure S3.

**Fig. S3 | *Aifm2*-deficiency in all T cells does not limit conventional T cell responses against bacteria but selectively diminishes Tregs during systemic *Listeria monocytogenes* infection**. (A) Experimental schematic for infection of wildtype versus mice with *Aifm2*-deficient T cells with $5x10^3$ CFU *Listeria monocytogenes*. (B) CFU analysis of spleen (left) and liver (right) for *Listeria monocytogenes* infection of wildtype versus mice with *Aifm2*-deficient T cells (n=4-5 mice/group). (C) Frequency of $Foxp3^+$ Tregs as a percentage of $CD4^+$(right) or total $CD45^+$ immune cells (left) in spleens of infected mice 6 days post infection. (D) Frequency of bulk $CD4^+$ Tconv cells as a total $CD45^+$ immune cells in spleens of infected mice 6 days post infection. (E) Frequency of bulk $CD8^+$ T cells as a total $CD45^+$ immune cells in spleens of infected mice 6 days post infection. (F) Frequency of Listeria-specific (SIIN-$H2Kb^+$) $CD8^+$ T cells in spleens of infected mice 6 days post infection. Data pooled from 2 experimental repeats (n=4-5 mice/group). For all plots, **P<0.05, **P<0.01, ***P<0.001 by student t-test, mean ± s.e.m.
